# Moving Beyond the ‘CAP’ of the Iceberg: Intrinsic Connectivity Networks in fMRI are Continuously Engaging and Overlapping

**DOI:** 10.1101/2021.06.23.449485

**Authors:** A. Iraji, A. Faghiri, Z. Fu, P. Kochunov, B. M. Adhikari, A. Belger, J.M. Ford, S. McEwen, D.H. Mathalon, G.D. Pearlson, S.G. Potkin, A. Preda, J.A. Turner, T.G.M. van Erp, C. Chang, V.D. Calhoun

## Abstract

Resting-state functional magnetic resonance imaging is currently the mainstay of functional neuroimaging and has allowed researchers to identify intrinsic connectivity networks (aka functional networks) at different spatial scales. However, little is known about the temporal profiles of these networks and whether it is best to model them as continuous phenomena in both space and time or, rather, as a set of temporally discrete events. Both categories have been supported by series of studies with promising findings. However, a critical question is whether focusing only on time points presumed to contain isolated neural events and disregarding the rest of the data is missing important information, potentially leading to misleading conclusions. In this work, we argue that brain networks identified within the spontaneous blood oxygenation level-dependent (BOLD) signal are not limited to temporally sparse burst moments and that these event present time points (EPTs) contain valuable but incomplete information about the underlying functional patterns.

We focus on the default mode and show evidence that is consistent with its continuous presence in the BOLD signal, including during the event absent time points (EATs), i.e., time points that exhibit minimum activity and are the least likely to contain an event. Moreover, our findings suggest that EPTs may not contain all the available information about their corresponding networks. We observe distinct default mode connectivity patterns obtained from all time points (AllTPs), EPTs, and EATs. We show evidence of robust relationships with schizophrenia symptoms that are both common and unique to each of the sets of time points (AllTPs, EPTs, EATs), likely related to transient patterns of connectivity. Together, these findings indicate the importance of leveraging the full temporal data in functional studies, including those using event-detection approaches.

## Introduction

Brain function results from the interaction among local and distributed brain areas. Thus, the temporal dependency among brain regions, commonly known as functional connectivity, is a widely-used tool for studying brain function (1). The advent of resting-state functional magnetic resonance imaging (rsfMRI) revealed the presence of strong temporal dependencies between functionally related brain regions (e.g., bilateral motor cortices) even without external stimulation (2). This led to a rapid growth in rsfMRI research and identification of spatial patterns of functionally connected regions termed intrinsic connectivity networks (ICNs), or functional networks, at different spatial scales from large-scale distributed networks to fine-grained, spatially local ones (3–5). Interestingly, spatial patterns obtained by applying independent component analysis (ICA) to spatial maps of thousands of different activation conditions derived from the BrainMap meta-analytic tool (https://www.brainmap.org) closely resemble large-scale networks (6), as do data-driven analyses of activation maps from task data (7), further supporting the functional relevance of functional networks. These approaches are implicitly built upon the notion of continuous information processing and interaction among brain regions.

In parallel to this work, another family of approaches, that we collectively called event detection approaches, was developed originally based upon a hypothesis that functional patterns and brain networks emerge from discrete, neural events (8, 9). Therefore, instead of studying static or dynamic functional patterns in the context of ongoing, continuous functional interactions, this family of approaches focuses on extracting discrete events to study static and dynamic functional patterns. The common procedure for these approaches is to first select a set of extreme time points (e.g., top 10%) as event present time points (EPTs) and use only these EPTs to obtain corresponding brain functional patterns (9, 10). There are also work that unified these two steps to directly obtain events and associated time points, for example, by applying clustering on all time points (11). These approaches assume that these event present time points alone are sufficient to estimate the temporal dependency and to derive the functional connectivity patterns observed in functional connectivity studies (12–14), and resting-state functional connectivity is driven by short-lived cofluctuation events (15). As such, event-based studies shift away from typical functional connectivity and mainly focus on capturing functional patterns as (co-)activation patterns of sparse EPTs using first-order statistics and signal amplitude rather than statistical dependence between them (9, 12, 16). Event detection approaches have shown direct correspondence between such spatial co-activation patterns and large-scale brain networks obtained from functional connectivity analysis (9, 11). They also demonstrated that different EPTs of a given node represent different co-activation patterns reflecting dynamic information of rsfMRI data (10, 17). However, these findings were limited to large-scale distributed brain networks with no established observations of fine-grained networks (ICNs) obtained from high-order ICA. This, in turn, may indicate that event detection approaches are limited to dominant large covarying functional patterns.

After years of using first-order statistics in event-based studies, which followed early work utilizing the temporal dependency between extreme time points to derive the functional connectivity (9, 12), recent studies revisited the idea of using extreme time points to study (dynamic) functional connectivity in the context of second and higher-order statistics and demonstrated the potential to provide additional insights into cognition and behavior (14, 15, 18).

These intriguing findings, along with the potential clinical and behavioral relevance of EPTs (19–21) and a number of supporting observations in animal studies (22–24), further buttress the assumption that rsfMRI signal may originate from brief, sporadically events rather than continuous dynamic functional interactions. This has led to a growing interest in studying brain function using momentarily sparse time points and related approaches (25–30).

However, an important question that has not yet been addressed is whether estimating results from only EPTs omits important information, potentially leading to misleading conclusions. While we do not refute the existence of discrete, neural events or even the possibility of large-scale networks are emerged from these spontaneous, infrequent events, we argue against the proposition that functional patterns estimated from the spontaneous blood oxygenation level-dependent (BOLD) signal manifest **only** in temporally sparse burst moments and these discrete EPTs contain incomplete information about the underlying functional patterns. This position can be supported by two specific observations. First, it has been shown that spiking activity propagates using both asynchronous and synchronous spiking (31). It is unlikely that sparse events alone can completely capture both of these modes. Second, the fact that connectivity estimators based on both amplitude (e.g., Pearson correlation) and phase (e.g., instantaneous phase synchrony) modulation that have been shown to contain meaningful information (4, 32) suggests that sparse events based only on amplitude in the space-time domain may give incomplete results.

We evaluated our premise by concentrating on the default mode as it is arguably the most detected/studied functional pattern in event-based studies and is commonly used to support the discrete nature of functional patterns in the spontaneous BOLD signal. We first asked whether the default mode pattern is present only in its so-called EPTs or whether it can also be identified from the event absent time points (EATs), i.e., time points with the least (almost zero) probability of being EPTs. Additionally, we explored whether the default mode EPTs contain similar functional connectivity information as the full data in the context of their associations with a diagnosis of schizophrenia and with schizophrenia symptoms. Finally, we asked whether the (usually ignored) EATs of the default mode could provide additional unique information about schizophrenia.

It should be mentioned that this study evaluates if default mode network continuously contributes to the BOLD signal, including the time points with the least probability of being EPTs. This is in contrast to earlier studies that assess the effect of scan length on the reliability of estimating static functional connectivity (33). Furthermore, while the term “co-activation pattern/map” is commonly used to describe spatial patterns identified by event-detection approaches, the term co-activation pattern (CAP) also refers to a specific category of event detection approaches (34), and the term co-activation has also been used in other contexts, such as describing task co-activation patterns across fMRI studies (35, 36). Therefore, we opted for the term activation spatial maps (ASMs) to describe the first-order activation patterns that are being evaluated in this study to avoid confusion. Furthermore, to simplify notation in the remainder of this paper, unless we explicitly indicate otherwise, the terms EATs and EPTs refer to the EATs and EPTs of the default mode.

## Materials and Methods

### Dataset and Inclusion Criteria

We used 3-Tesla rsfMRI data, comprising 477 typical controls and 350 individuals with schizophrenia selected from three datasets (Table 1) with different data acquisition parameters, including Functional Imaging Biomedical Informatics Research Network (FBIRN) (37), Center for Biomedical Research Excellence (COBRE) (38), and Maryland Psychiatric Research Center (MPRC) (39).

**Table 1.** Demographic information of Subjects studied. FBIRN: Functional Imaging Biomedical Informatics Research Network, COBRE: Center for Biomedical Research Excellence, and MPRC: Maryland Psychiatric Research Center.

| Dataset | Diagnostic (#) | Sex (#) | Age (years) |  |
| --- | --- | --- | --- | --- |
| | | | mean ± sd | median/range |
| **FBIRN** | *Control group (160)* | Male (115) | 37.26 ± 10.71 | 39/(19-59) |
| | | Female (45) | 36.47 ± 11.33 | 33/(19-58) |
| | *Schizophrenia group (150)* | Male (114) | 38.74 ± 11.78 | 40/(18-62) |
| | | Female (36) | 39.06 ± 11.40 | 36/(21-57) |
| **COBRE** | *Control group (79)* | Male (55) | 39.07 ± 12.43 | 38/(18-65) |
| | | Female (24) | 34.92 ± 10.23 | 34/(18-58) |
| | *Schizophrenia group (50)* | Male (42) | 37.43 ± 15.05 | 32.5/(19-64) |
| | | Female (8) | 43.25 ± 12.78 | 40/(31-65) |
| **MPRC** | *Control group (238)* | Male (94) | 38.72 ± 13.63 | 40/(12-68) |
| | | Female (144) | 41.22 ± 16.06 | 44/(10-79) |
| | *Schizophrenia group (150)* | Male (98) | 35.57 ± 13.18 | 32/(13-63) |
| | | Female (52) | 44.60 ± 13.87 | 47/(13-63) |

The FBIRN includes rsfMRI data collected at six sites using Siemens 3-Tesla Tim Trio scanners and one site using a General Electric 3-Tesla Discovery MR750 scanner. All sites use the same following parameters: a standard gradient echo-planar imaging sequence, repetition time/echo time (TR/TE) = 2000/30 ms, voxel spacing size = 3.4375 × 3.4375 × 4 mm, slice gap = 1 mm, field of view = 220 × 220 mm, and a total of 162 volume. The COBRE dataset was collected at one site using a Siemens 3-Tesla TIM Trio scanner and a standard echo-planar imaging sequence with TR/TE = 2000/29 ms, voxel spacing size = 3.75 × 3.75 × 4.5 mm, slice gap = 1.05 mm, field of view = 240 × 240 mm, and a total of 149 volumes. Finally, the MPRC dataset was collected in three sites: (1) Siemens 3-Tesla Allegra scanner using a standard echo-planar imaging sequence with TR/TE = 2000/27 ms, voxel spacing size = 3.44 × 3.44 × 4 mm, FOV = 220 × 220 mm, and 150 volumes; (2) Siemens 3-Tesla Trio scanner using a standard echo-planar imaging sequence with TR/TE = 2210/30 ms, voxel spacing size = 3.44 × 3.44 × 4 mm, FOV = 220 × 220 mm, and 140 volumes; and (3) Siemens 3-Tesla Tim Trio scanner using a standard echo-planar imaging sequence with TR/TE = 2000/30 ms, voxel spacing size = 1.72 × 1.72 × 4 mm, field of view = 220 × 220 mm, and 444 volumes.

The inclusion criteria include 1) the number of time points (volumes) of the BOLD time series larger than 100 (the minimum number of time points is 135), 2) head motion transition less than 3° rotations and 3 mm translations in every direction, 3) mean framewise displacement less than 0.25, 4) high-quality registration to an echo-planar imaging template, and 5) spatial overlap between individual mask and group mask above 80%.

### Preprocessing

The rsfMRI data were preprocessed primarily using the statistical parametric mapping (SPM12, http://www.fil.ion.ucl.ac.uk/spm/) toolbox. The first five volumes were discarded for magnetization equilibrium purposes. Rigid body motion correction was performed to correct subject head motion during the scan. Slice-timing correction was applied to correct timing differences (temporal misalignment) in slice acquisition. The data of each subject were warped into a Montreal Neurological Institute (MNI) echo-planar imaging template and resampled to 3 mm^3^ isotropic voxels. Next, the data were spatially smoothed using a Gaussian kernel with a 6 mm full width at half-maximum. Considering ICA separates white matter and cerebrospinal fluid signals into independent components, we do not regress them out during preprocessing (thus following standard practice for ICA analyses).

Because the fMRI signal has a low signal-to-noise ratio, the detection of EPTs can be sensitive to noise. For example, noise can change the time points that survive thresholding and alter the time points that manifest as local maxima. To address this, we performed the following cleaning steps on the time course of each voxel. Voxel time courses were detrended by removing linear, quadratic, and cubic trends. We regressed out six motion realignment parameters and their derivatives. Outliers were detected based on the median absolute deviation, spike threshold (c_1_) = 2.5 (40), and replaced with the best estimate using a third-order spline fit to the clean portions of the time courses. Bandpass filtering was applied using a fifth-order Butterworth filter with a cutoff frequency of 0.01 Hz-0.15 Hz. We also evaluated if the cleaning procedure drives the findings by repeating the analysis using uncleaned data.

### Extracting Default Mode Time Course

We used several procedures to obtain the time series associated with the default mode network to illustrate that our findings are not the result of a selected node or time courses. Similar to most event-based studies, we first used a region of interest (ROI) in the posterior cingulate cortex area as a node for the default mode. For this purpose, we used the two most commonly used ROIs including a 6 × 6 × 6 mm^3^ cube centered at (x = 0, y = −53, z = 26) (10) and a sphere with 3mm-radius, centered at (x = −6, y = −58, z = 28)) (41) in MNI coordinates. We called these two Node_Seed1_ and Node_Seed2_. We defined an additional node (Node_Meta_) using a term-based meta-analysis for the term “default mode” in Neurosynth (https://www.neurosynth.org/) (42). The default mode associated map was registered to the study common space, and the top 27 voxels were selected as Node_Meta_ for the default mode. The Node_Meta_ includes a cluster of size 20 voxels in the posterior cingulate cortex and a cluster of size seven voxels in the left angular gyrus. For these three nodes, the average time course within each node was used as the representation of the default mode time course to identify time points with the highest and lowest contributions of the default mode.

While the default procedure for existing event-based approaches is to use anatomically fixed nodes to estimate EPTs, one may argue our findings are the result of using predefined anatomically fixed nodes for all individuals, which could result in inaccurate estimation of the default mode time course. As such, we also utilized ICA to obtain subject-specific default mode maps and associated time courses (Node_ICA_) (43, 44). ICA was performed using the GIFT v4.0c software package (https://trendscenter.org/software/gift/) (45). We applied a low model order (model order = 20) group-level spatial ICA to obtain large-scale brain networks (46), including the default mode. We applied variance normalization (z-score) on voxel time courses and computed subject-level spatial principal components analysis (PCA) to retain maximum subject-level variance (greater than 99%). Subject-level principal components were then concatenated together across the time dimension, and group-level spatial PCA was applied to the concatenated subject-level principal components. The 20 group-level principal components that explained the maximum variance were selected as the input for the Infomax algorithm to calculate 20 group independent components. The Infomax ICA algorithm was run 20 times, and only components with (ICASSO cluster quality index > 0.8) were used to identify networks. Next, we used the group component as a reference in a spatially constrained ICA (called group information guided ICA or GIG-ICA) to calculate subject-specific default mode and its associated time courses (47).

### Detecting Event Present and Absent Time Points (EPTs and EATs) of the Default Mode

The main idea behind event-based studies is that functional connectivity patterns, such as the default mode, are derived from sparse, abrupt neuronal events which manifest as momentarily changes in the amplitude (or other characteristics) of the BOLD signal of associated regions. Thus, the key step for event-based studies is to identify a set of time points with the largest association to a predefined node (i.e., EPTs). Here, to evaluate our hypothesis, we also focus on the set of time points with the weakest association to the predefined node (i.e., EATs), which are at the opposite extreme, the time points with the minimum chance of event occurrence. Event-based studies use different techniques to detect EPTs from the time series of a given node, which are suggested to lead to similar results (13). The most common techniques include 1) choosing the time points with maximum amplitudes (e.g., top 10 percent or top 20 percent time points), 2) choosing time points that pass a threshold value (e.g., above one standard deviation of the time series), 3) selecting time points which are the local maxima/minima of the time series, and 4) deconvolving the BOLD signal using hemodynamic models, which resolve into a set of sparse EPTs.

Focusing on the default mode, our main objective here is not to identify the default mode and its EPTs, but rather to assess the possibility of estimating the default mode from its assumed EATs, and secondarily whether we are losing information if we focus only on the EPTs. In other words, we first aim to evaluate if the default mode can be retrieved by only using EATs. We can use any event detection techniques and assign time points not selected as EPTs as EATs, but here we chose a more extreme approach and defined EATs as time points that are the least likely EPTs of the default mode. For this purpose, we selected the 20 time points (equal to the ICA model order) with the minimum amplitude of a given node time series as EATs (**Figure 1**). In the same manner, the 20 time points with the largest amplitude are considered to be the EPTs (**Figure 1**). We also evaluate the finding by choosing the 20 time points with the highest absolute value.

**Figure 1.**
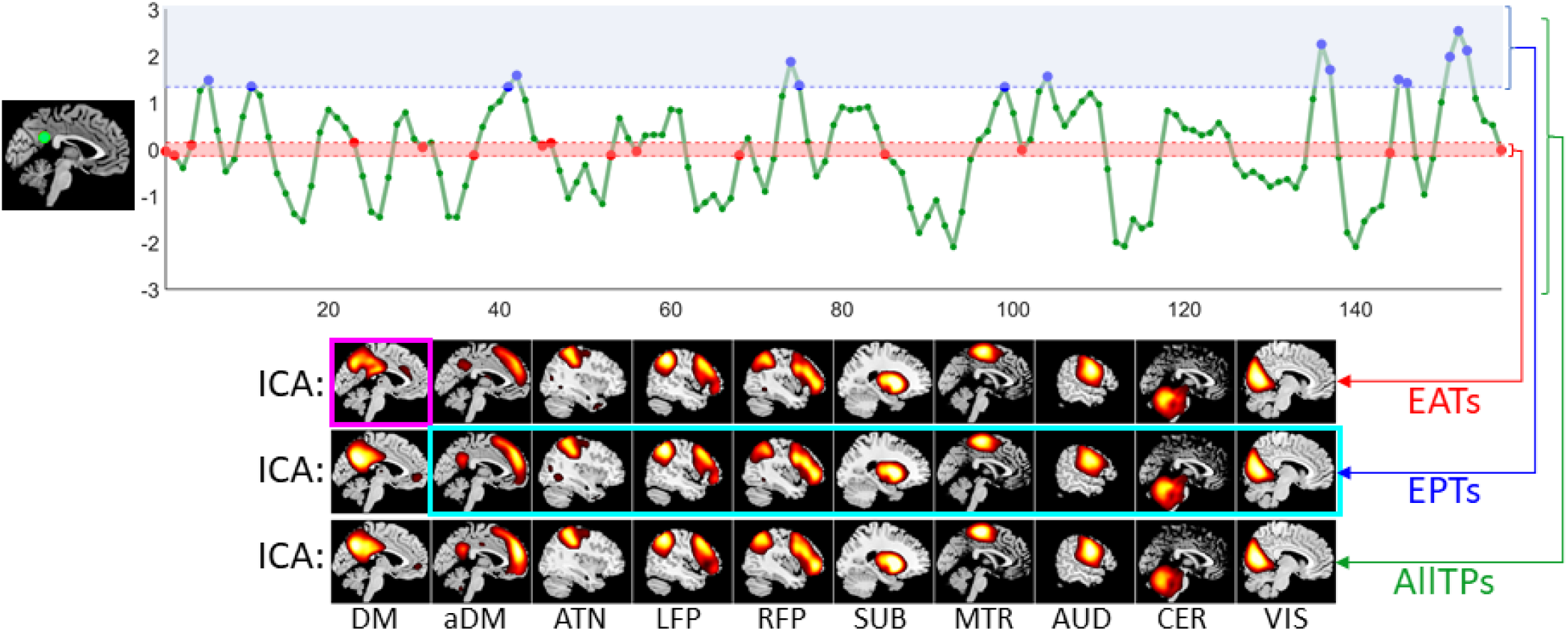
Schematic of the analysis pipeline. First, we select one of the four default mode nodes and use its time course to identify event present time points (EPTs) and event absent time points (EATs) for the default mode. The 20 time points with the largest amplitude are labeled as the EPTs (blue dots), while EATs are the 20 time points with minimum amplitudes (red dots). We selected this time point selection technique to minimize the likelihood of the presence of an event in EATs. We also repeated the analysis by using the amplitude of absolute value (as some studies use both positive and negative extremes as EPTs). We compared EPTs and EATs with the functional patterns obtain using all time points (AllTPs, green dots). We evaluated the presence of functional patterns in EATs, EPTs, and AllTPs of default mode (DM) using different tools, including high-order statistics using independent component analysis (ICA), presented in the figure, second-order statistics using pair-wise Pearson correlation, and first-order statistics using (weighted-) activation spatial maps (ASMs). We were specifically interested in whether (1) the DM is present within the EATs (magenta box) and (2) EPTs of the DM also include information about the non-DM networks (cyan box). aDM: anterior default mode network (as opposed to classical (posterior) DM), ATN: attention network, LFP/RFP: left/right frontoparietal network, SUB: subcortical network, MTR: Somatomotor network, AUD: auditory network, CER: cerebellar network, and VIS: visual network.

In addition, we used EPTs to investigate if other large-scale networks can be obtained by applying ICA to the EPTs of the default mode (**Figure 1**). In other words, we evaluated if the default mode EPTs are only occupied or mainly dominated by default mode or other networks similarly present and contribute to these time points. While some recent event-based works consider the possibility of the coexistence of a few functional patterns and allow the temporal overlap between their events (17, 48), event-based studies commonly assume the activation peaks of different networks do not coincide together. Here, we argue for the continuous presence of all networks including EPTs of other networks and assess this by focusing on EPTs of the default mode. The presence of other networks in the EPTs of the default mode (and also EATs and AllTPs) also suggests the continuous presence of overlapping networks.

### Evaluating Event Present and Absent Time Points (EPTs and EATs)

We used the following procedures to evaluate the presence of the large-scale networks in the EATs (EPTs) of the default mode estimated using the various nodes mentioned earlier. First, we applied ICA to the EATs (EPTs) of the default mode to evaluate if we can obtain brain networks from these time points. We used ICA with the same parameters as explained above. However, instead of using all time points, we only used the EATs (EPTs) of each subject for this procedure. To further clarify, ICA analysis has been used for different purposes. Once, we use ICA to estimate subject-specific default mode node which is called Node_ICA_. The other time, we applied ICA analysis to extract ICNs (including the default mode network) from different portions of data (EPTs/EATs). Moving forward, whenever we talk about ICA results we are referring to the second one (i.e., running ICA on EPTs/EATs). We also calculated whole-brain functional connectivity maps by calculating Pearson correlation between a node time series and all brain voxels using only EATs (EPTs). The main objective of this procedure was to determine if the default mode can be obtained from EATs using linear second-order statistics or if high-order statistics (for example, using ICA) are required to extract the network.

Finally, we calculated whole-brain ASMs to evaluate if the co-activation pattern of the default mode can be estimated from EATs in addition to EPTs. First, we calculated the weighted-ASM (wASM) (Eq. 1), where weight at each time point, *w(t)*, is node signal amplitude and *X(t)* is the preprocessed BOLD signal at a given time point.

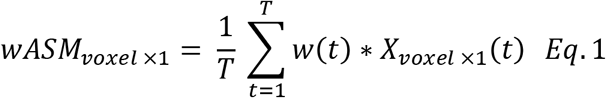

We also calculated ASM by averaging EATs (EPTs) while correcting the sign of node signal at a given time point (Eq. 2). The sign correction is needed to prevent positive and negative values from canceling each other. When EPTs are obtained using the amplitude of a given node time course, EPTs are expected to have positive values, and Eq. 2 is equal to the simple averaging. However, many studies use the maximum absolute values of a node time course or L^1^-norm (or L^2^-norm) of time series obtained from several node amplitudes at any given time, which requires sign correction.

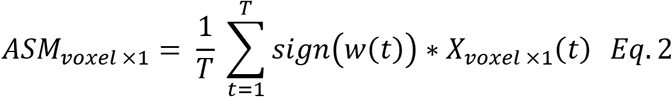

For all analyses, we use spatial correlation to quantify the similarities in the spatial maps. To ensure that the effect of other networks is not driving similarity values, we first regressed out the contribution of other networks.

### Recursive ICA

When we use GIG-ICA to obtain a subject-specific node for the default mode (Node_ICA_), EATs are estimated from subject-specific default mode time courses. Thereby, the EATs are largely dominated by other networks, and typically it is assumed there would be no significant presence of default mode within these timepoints. However, we hypothesized that even in this scenario, the default mode network is present and can be recovered. To evaluate our hypothesis, we introduce an approach called recursive ICA, which is simply performing ICA on the residual of the previous ICA. For this purpose, we first estimate ICA components from the EATs and obtained the stable components. Then, we regress out these components from each subject EATs and estimate components from the residual. Recursive ICA is similar to Snowball ICA approach (49) in which information and ICNs are collected iteratively by using the recursive ICA subtraction approach.

### Event Present and Event Absent Time Point (EPT and EAT) Signatures in Schizophrenia

We next investigated whether default mode EPTs carry all information about the default mode or if instead different temporal portions, EPTs, EATs, and all time points (AllTPs), contain distinct and potentially complementary information about the default mode. We used Node_ICA_ as it uses subject-specific default mode information to provide the best estimation of the default mode EPTs and EATs from data itself.

We first evaluated the association between a schizophrenia symptom score and the default mode functional network connectivity (DM-FNC). Because different symptom scores, the Positive and Negative Syndrome Scale (PANSS) and the Brief Psychiatric Rating Scale (BPRS), were recorded for different individuals with schizophrenia, we used a prior conversion algorithm obtained from 3767 individuals to convert BPRS total scores to PANSS total scores (50) and used PANSS total score in our analysis. We calculated FNC between the default mode network and nine other large-scale networks for EPTs, EATs, and AllTPs (aka, static DM-FNC). We regressed out age, gender, site, and mean framewise displacement from each FNC. For each approach, i.e., AllTPs, EPTs, and EATs, we used nine DM-FNCs as the independent variables and the total PANSS score as the dependent variable. We applied the least absolute shrinkage and selection operator (LASSO) (51) with ten-fold cross-validations (CV = 10) and 50 Monte Carlo repetitions for cross-validation to identify the symptom-related DM-FNC features. We chose the model and the regularization coefficient (lambda) that resulted in the minimum cross-validated mean squared error (MSE). The DM-FNCs with nonzero coefficients are the most relevant attributes, among the independent variables in the model, to best describe the PANSS score in our dataset. If all coefficients are zero for a model, it means none of the FNC pairs significantly (i.e., beyond constant value) relate to the symptom severity score. Next, we evaluated if symptom-related DM-FNCs (DM-FNCs with nonzero coefficients, if there are any), obtained using individuals with schizophrenia, can also differentiate between individuals with schizophrenia and typical controls using a logistic regression model. Similar to the previous analysis, the logistic regression was applied after regressing out age, gender, site, and mean framewise displacement covariates.

## Results

### The Default Mode Never Rests

The results for using the amplitude of Node_Seed1_ time series to obtain the default mode from EATs and EPTs are presented in **Figure 2**. The ICA results (i.e., running ICA on EATs obtained using Node_Seed1_) show that the default mode network can be estimated well from just the EATs. The spatial similarities between subject-level and group-level default mode, while controlling for the effect of other networks, are 0.674 ± 0.053 (mean ± SD) for AllTPs, 0.483 ± 0.072 for EPTs, and 0.366 ± 0.083 for EATs. If we consider this criterion as the reproducibility measurement, results from AllTPs outperform EPTs, suggesting the potential benefit of using all time points. The result for EATs shows lower spatial similarity at the subject-level (group-level results are comparable); however, the similarity is well above chance. We also performed an additional test by removing local peaks from EATs (i.e., EATs are less than 20 time points) and applying ICA on the remaining EATs, which shows the presence of the default mode network.

**Figure 2.**
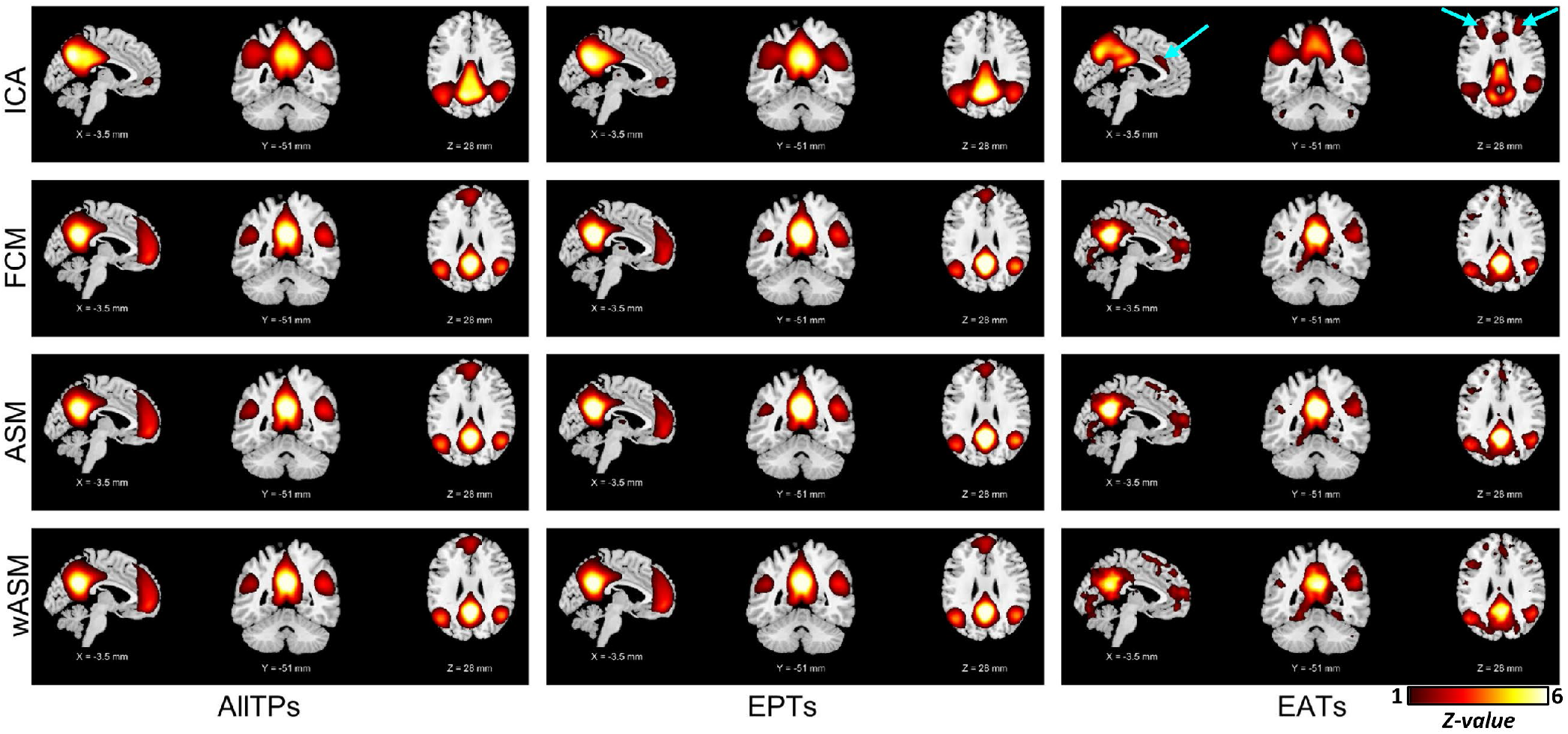
NodeSeed1 default mode functional patterns. The plots are z-maps (mean equals to zero and standard deviation equals to one), thresholded at *z-*value = 1. The z-score colorbar range is [1 - 6] for all plots. The first column shows the functional patterns obtained using all time points (AllTPs) of individuals. The second column represents the results obtained by applying analysis on the event present time points (EPTs), and the third column is the functional patterns obtained using only event absent time points (EATs). The first row is showing the ICA results. The top left shows the default mode network calculated by applying ICA on the whole dataset (using all time points). The default mode network was successfully obtained for both EPTs and EATs. For EATs, NodeSeed1, which is in the posterior cingulate area, does not strongly contribute to the default mode network (see the ‘hole’ in seed location in the top right panel. However, even in EATs, the NodeSeed1 connectivity with the default mode regions is stronger than its connectivity with the rest of the brain as shown in the functional connectivity map (FCM) of NodeSeed1 obtained using pair-wise Pearson correlation (the second row). In other words, this result suggests the default mode network exists strongly at EAT time points, and NodeSeed1 is functionally connected to it. Activation spatial maps (ASMs) and weighted ASMs (wASMs) show very similar co-activation patterns for AllTPs, ETPs, and EATs consist of default mode regions. Indeed, the EATs maps are noisier because the small amplitude of NodeSeed1 time course in EATs making it more susceptible to noise. The arrows in cyan color show regions commonly contribute to the executive control network.

As expected, the spatial patterns of the default mode in EATs demonstrated meaningful differences. For example, the location of Node_Seed1_ itself does not strongly contribute to the default mode network in these EATs, which partially explains the lower spatial similarity in EATs. In other words, the default mode exists strongly in Node_Seed1_ EATs, but the contribution of Node_Seed1_ to the default mode in EATs is not as strong as the rest of the default mode regions. The fact that the default mode can still be estimated from the EATs, where it should be least likely to be detected, provides evidence that intrinsic networks are best modeled as continuous in both space and time, rather than as discrete events.

We next studied the connectivity profile of Node_Seed1_ in the EATs to evaluate if Node_Seed1_ is completely dissociated from the default mode or, despite not being a part of the default mode network dominant pattern, Node_Seed1_ maintains its connection to the default mode regions. Interestingly, even in the EATs, Node_Seed1_ is significantly connected to the default mode regions, and the default mode connections are the dominant functional connectivity of Node_Seed1_ even in its EATs (**Figure 2**). The activity patterns further supported the functional connectivity results in that both ASMs and weighted ASMs (wASMs) show co-activation patterns resembling the default mode network for AllTPs, ETPs, and EATs. The connectivity and co-activity patterns of Node_Seed1_ in EATs are somewhat noisier because of the small amplitude and fluctuations of Node_Seed1_ time series in these time points; however, the default mode pattern is still clearly present. The spatial similarities between subject-level and group-level default mode for these analyses can be found in Supplementary 1.

We also calculated the spatial similarities of EPTs (and EATs) with AllTPs (Table 2) at both subject-level and group-level. For both EPTs and EATs, the spatial similarity is highly significant. EPTs shows higher similarity compared to EATs, which is expected as EPTs contribute more than EATs to the spatial maps obtained from AllTPs. The lower spatial similarity in EATs is also derived by a different topology of the default mode in EATs, which is related to the dynamic nature of the brain, including the lower contribution of the posterior cingulate cortex to the default mode, which significantly contributes to EPTs. Another example of different topologies can be seen in ICA analysis (i.e., running ICA on EATs and EPTs). In the EAT ICA analysis, many regions of the executive control network (identified as the cyan arrows in **Figure 2**) have merged with the default mode component, suggesting these two networks are dynamically integrated with one another into a single network. In contrast, these two networks are identified separately in EPTs and AllTPs analysis.

**Table 2.** The spatial similarity between AllTPs with EPTs and EATs using NodeSeed1: “subject-level mean ± subject-level standard deviation (group-level)”.

| Node<sub>Seed1</sub> | AllTPs - EPTs | AllTPs - EATs |
| --- | --- | --- |
| ICA | $0.648 \pm 0.089$ (0.972) | $0.371 \pm 0.084$ (0.840) |
| FCM | $0.520 \pm 0.178$ (0.927) | $0.513 \pm 0.197$ (0.902) |
| ASM | $0.576 \pm 0.164$ (0.733) | $0.499 \pm 0.188$ (0.891) |
| wASM | $0.666 \pm 0.150$ (0.865) | $0.504 \pm 0.211$ (0.895) |

Performing analysis using the absolute value of Node_Seed1_ time series resulted in similar findings as using the signed amplitude of Node_Seed1_ time series (Supplementary 2). The findings were replicated when other nodes were used to identify EPTs and EATs. **Figure 3** (and Supplementary 3 and 4) shows the results for Node_Seed2_ and Node_Meta_ when the absolute value (amplitude) of their time courses were used to extract EATs and EPTs. These results are similar to those for Node_Seed1_, suggesting our findings are not the results of specific node or event selection procedures. Our analysis also shows that the spatial similarity of the default mode is well above chance for all of the nodes. For instance, the subject-level similarities between functional connectivity maps obtained from Node_Seed1_ and Node_Meta_ are 0.909 ± 0.076 (mean ± SD) for AllTPs, 0.689 ± 0.221 for EPTs, and 0.460 ± 0.244 for EATs, which indicate more robustness toward node selection for AllTPs compared to EPTs and EATs in static analysis. The results are the same for other analyses, including ASM and wASM (Supplementary 5). It should be noted that while the sensitivity to node selection could be related to the regions’ functional specificity and brain dynamics in addition to lower SNR, it has an undesirable effect on the reproducibility of static analysis particularly in the EATs which show lower subject-level similarity. Thus, caution should be taken when using EATs to evaluate within network connectivity patterns, especially in clinical studies.

**Figure 3.**
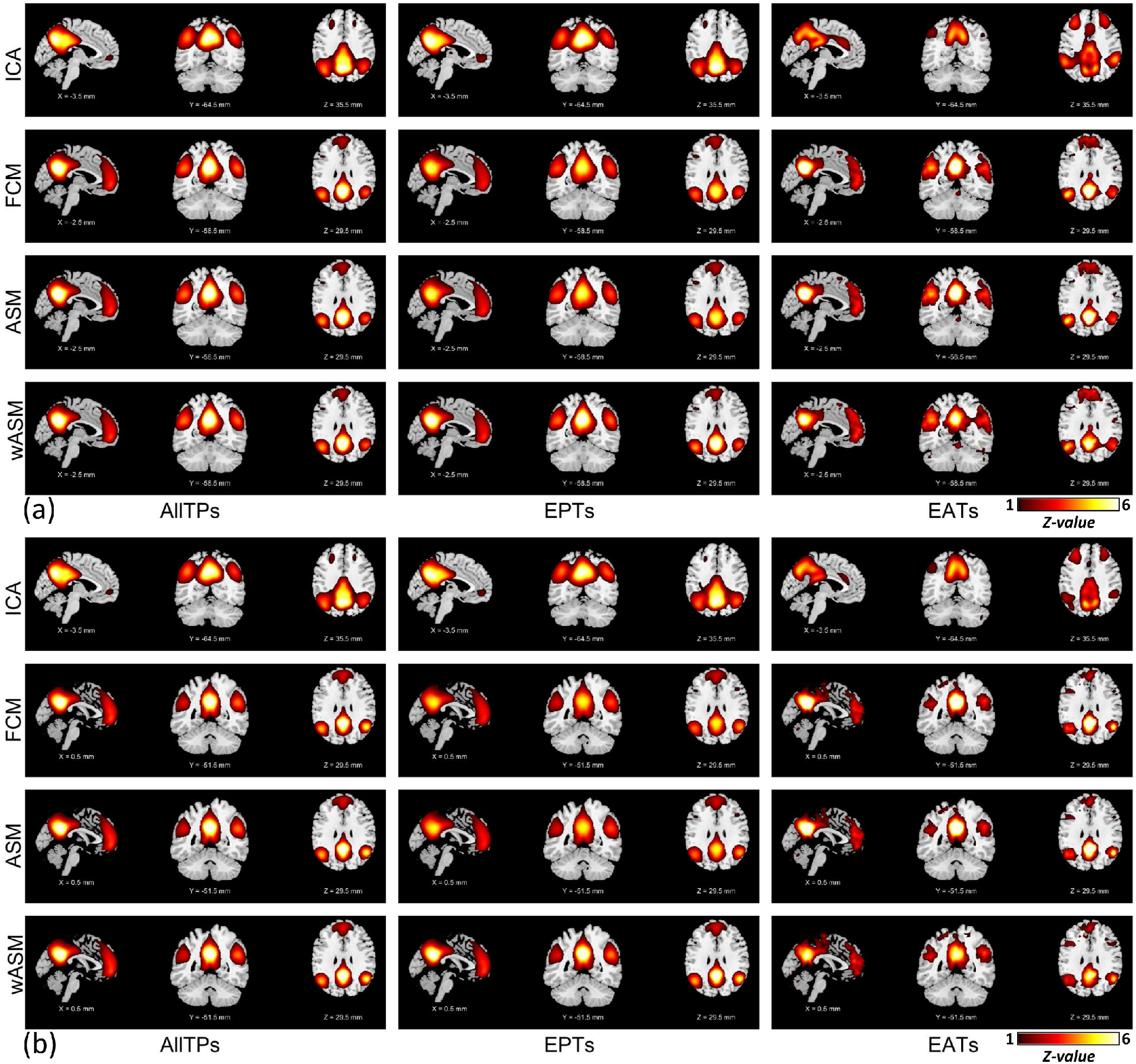
The default mode functional patterns were obtained using the absolute value of NodeSeed2 (a) and NodeMeta (b) time courses. The plots are z-maps (mean equals to zero and standard deviation equals to one), thresholded at z-value = 1. The z-score colorbar range is [1 - 6] for all plots. The first, second, and third columns illustrate the results obtained using all time points (AllTPs), using only the event present time points (EPTs), and using only event absent time points (EATs), respectively. NodeSeed2 and NodeMeta results reproduce the findings obtained from NodeSeed1 (see Figure 1).

Next, we evaluated the presence of the default mode in EATs (and EPTs) identified from the time course of the subject-specific default mode network (i.e., Node_ICA_). As we predicted, because these EATs were obtained from the same subject’s data, the default mode network activity was weak and concealed by the strong activity of other sources (such as other networks and artifacts). By using recursive ICA and removing the contribution of dominant components from EATs first, we successfully estimated the default mode network from the EATs alone (**Figure 4**). We also observed significant functional connectivity and co-activation in EATs between the time course of the default mode network and other regions involved in the default mode (**Figure 4**). The result confirmed the previous findings using other nodes, suggesting the default mode and functional connectivity between associated regions consistently present in spontaneous BOLD signals. Results were also consistent when we did not perform cleaning steps on the voxel time course.

**Figure 4.**
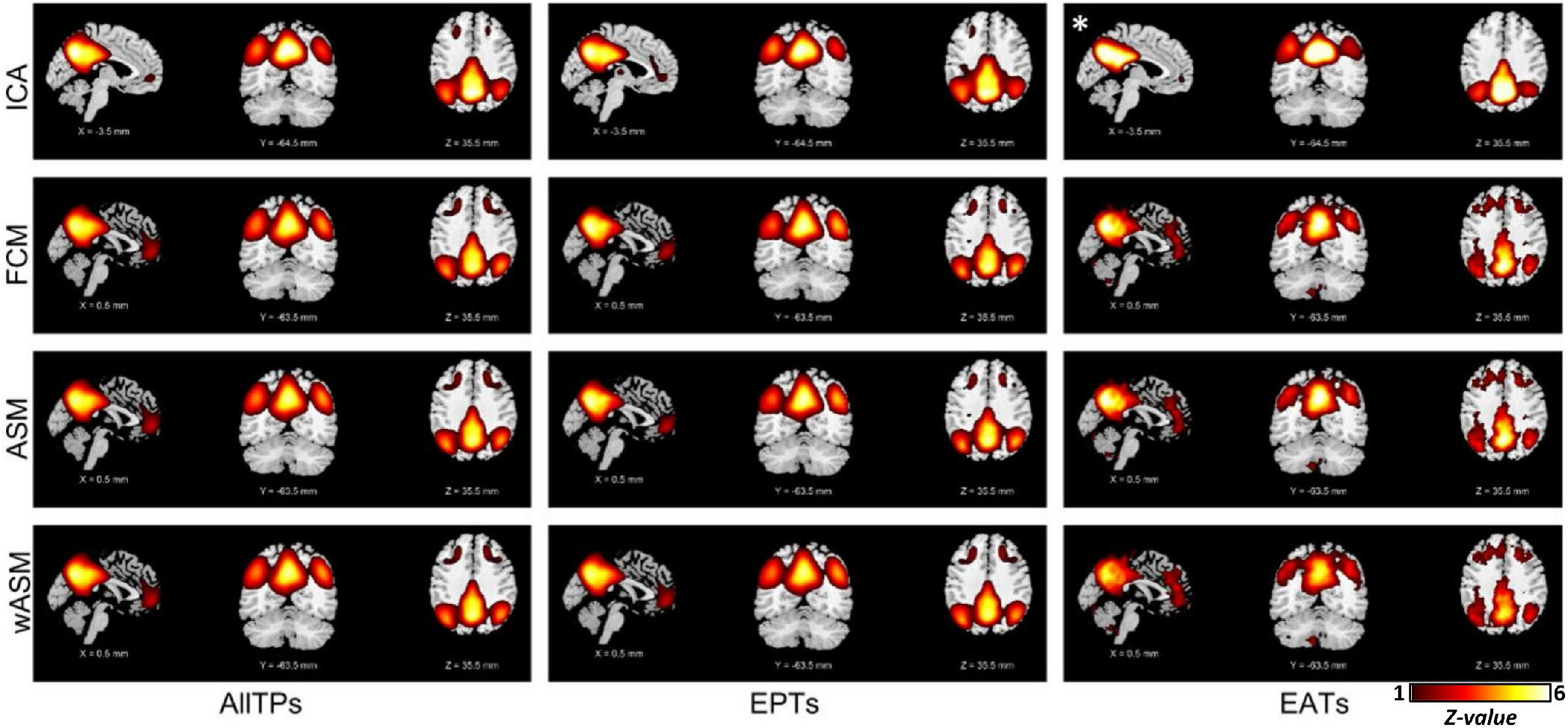
The default mode functional patterns were obtained using the amplitude of NodeICA. NodeICA is subject-specific default mode network obtain using group-information guided independent component analysis (GIG-ICA) (47). The plots are z-maps (mean equals to zero and standard deviation equals to one), thresholded at z-value = 1. The z-score colorbar range is [1 - 6] for all plots. The first, second, and third columns illustrate the results obtained using all time points (AllTPs), using only the event present time points (EPTs), and using only event absent time points (EATs), respectively. * indicates the default mode network obtain using recursive ICA.

In a supplementary analysis, we calculate the contribution of the default mode across time for 50 randomly selected subjects. The contribution values are the coefficient (beta) from linear regression of the default mode onto the fMRI data. **Figure 5** shows the sorted contribution of the default mode with no sudden drop in its contribution in any of these subjects. The results are the same for other studied networks, including the visual and somatomotor networks (Supplementary 6). We additionally evaluate and observe that the visual and somatomotor networks can be obtained from their own EATs (Supplementary 7).

**Figure 5.**
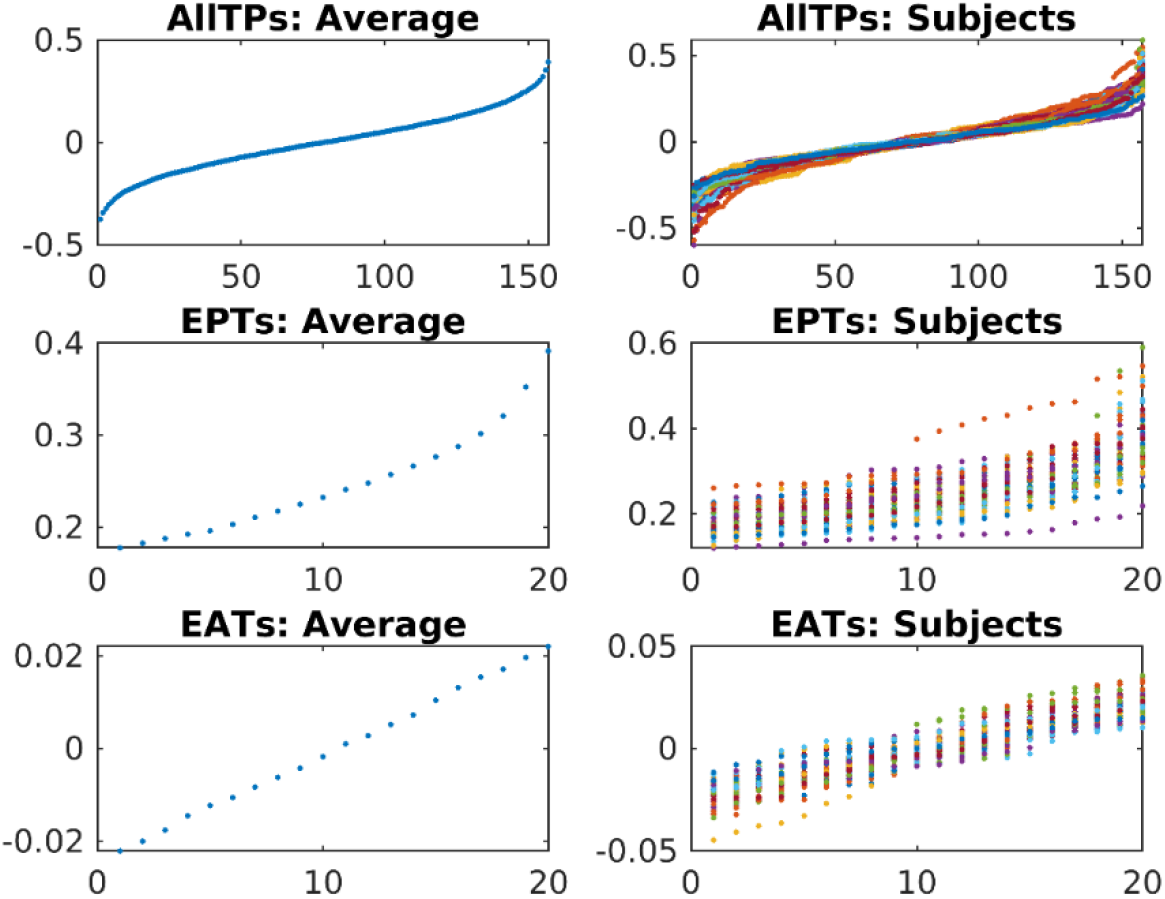
The sorted contribution of the default mode to the BOLD signal over time. The left column shows the average contribution across 50 randomly selected subjects, and the figures in the right column show contributions for each of the 50 individuals. The first row represents sorted contributions across all time points. The second and third columns show contributions to EPTs and EATs obtained using NodeICA. The results show smooth changes in the default mode contribution to time points with no sudden change that can explain changes related to before and after the occurrence of an event.

### The Default Mode Event Time Points are also Occupied by Other Networks

Next, we evaluated the presence of large-scale networks obtained in static analysis using all time points in EPTs and EATs. For all investigated nodes (i.e., Node_Seed1_, Node_Seed2_, Node_Meta_, and Node_ICA_), we successfully obtained the large-scale networks for both EPTs and EATs, suggesting that the large-scale networks present in the data regardless of the strength of the default mode and associated regions. **Figure 6** shows the large-scale networks obtained by applying ICA on the amplitude value of Node_Meta_ (i.e., Neurosynth default mode associated map) time course. The large-scale networks for other nodes and conditions can be found in Supplementary 8 and 9.

**Figure 6.**
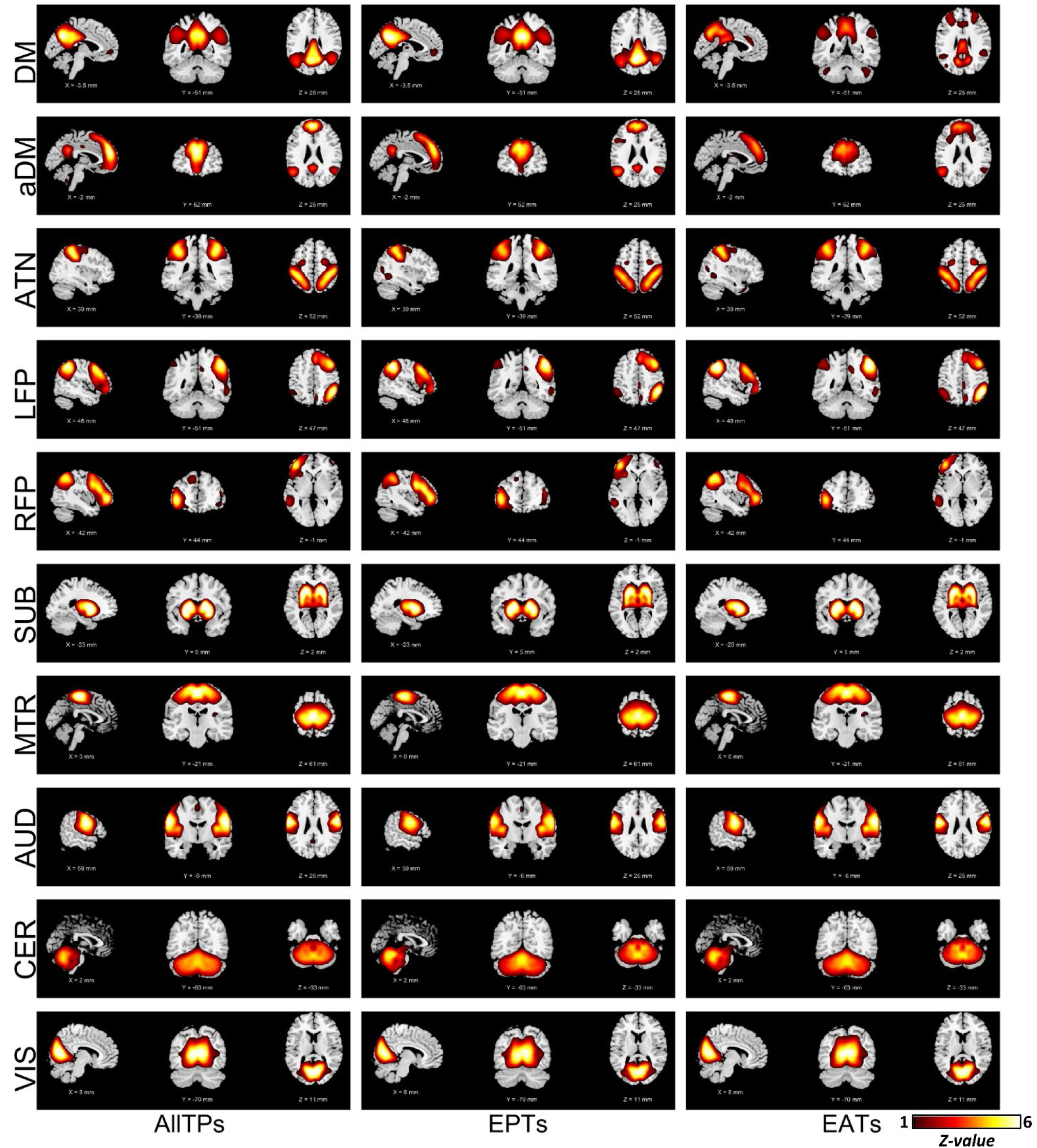
The large-scale networks calculate using the amplitude value of NodeMeta time courses. The plots are z-maps (mean equals to zero and standard deviation equals to one), thresholded at z-value = 1. The z-score colorbar range is [1 - 6] for all plots. The first, second, and third columns illustrate the results obtained using all time points (AllTPs), using only the event present time points (EPTs), and using only event absent time points (EATs), respectively. This result shows EPTs are not solely dominated by the default mode network, and the large-scale networks are equally present in EPTs. We also retrieve large-scale networks from EATs, suggesting the networks consistently present in the BOLD signal. DM: default mode network (also known as posterior DM). aDM: anterior default mode network (as opposed to classical (posterior) DM), ATN: attention network, LFP/RFP: left/right frontoparietal network, SUB: subcortical network, MTR: Somatomotor network, AUD: auditory network, CER: cerebellar network, and VIS: visual network.

These findings are important because we show other networks present in the EPTs of the default mode, providing evidence that the activation peaks of the default mode do not only contain the default mode but also similarly contain other networks. In other words, by showing networks exist in EPTs of the default mode, we provide evidence of the presence of brain networks beyond isolated events, complementary evidence of continuous present coexistence networks.

### Event Present and Event Absent Time Point (EPT and EAT) Unique Signatures in Schizophrenia

We next studied the DM-FNC obtained from different portions (EPTs, EATs, and AllTPs) of data in the context of their associations with schizophrenia and schizophrenia symptoms. We observed significant associations with PANSS score for different portions (for EATs: *|r|/p* = 0.191/0.002; for EPTs: *|r|/p =* 0.143/0.018; and for AllTPs: *|r|/p* = 0.125/0.038). Post-hoc analysis shows that the weights of symptom-related DM-FNCs (1) are almost identical across LASSO repetitions and (2) did not show significant differences for regression model applied across different subsets of data. Moreover, the model obtained for each time portion is not significantly associated with PANSS in other time portions.

**Figure 7** shows the minimum cross-validated mean squared error results for generalized linear models obtained using LASSO with ten-fold cross-validations and 50 repetitions, where the response variable (dependent variable) is the total PANSS score and the predictor variables (independent variables) are nine DM-FNCs obtained using AllTPs, EPTs, or EATs. The lines in **Figure 7** show the symptom-related DM-FNCs for AllTPs model in red, EPTs model in blue, and EATs model in green, and the width of each line represents the magnitude of the coefficients. The connection between the default mode and the cerebellum (DM-CER) contributes to all three models, suggesting its steady effect over time. However, it has much smaller coefficient in EPTs (|b|= 0.61) compared to AllTPs (|b| = 2.14) and EATs (|b| = 2.81). We also observed unique symptom-related patterns across models. For example, the default mode and the auditory (DM-AUD) connection contributes only to AllTPs model (|b| = 0.58), the default mode and the right frontoparietal (DM-RFP) connection contributes only to EPTs model (|b| = 0.28), and the default mode and the left frontoparietal (DM-LFP) connection contributes only to EATs model (|b| = 4.54).

**Figure 7.**
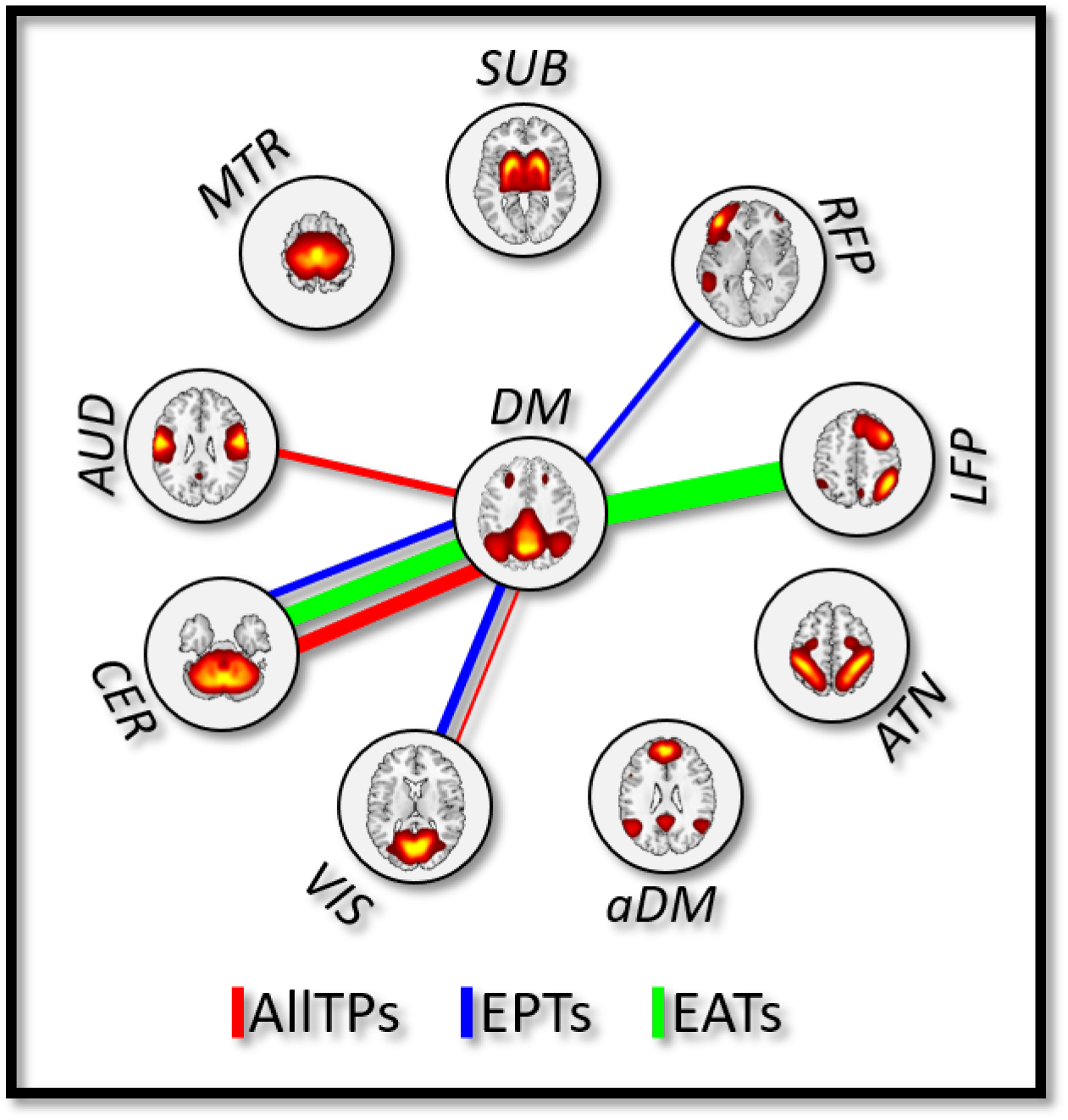
The association between schizophrenia symptoms and the default mode functional network connectivity (DM-FNC). The lines represent the coefficients of the generalized linear model obtained using the least absolute shrinkage and selection operator (LASSO) with ten cross-validations (CV = 10) and 50 Monte Carlo repetitions for the model with the minimum cross-validated mean squared error (MSE), where the response variable is the total of positive and negative syndrome scale (PANSS) scores and the predictor variables are nine default mode-FNC pairs estimated using all time points (AllTPs) in red, using the default mode event present time points (EPTs) in blue, and using the default mode event absent time points (EATs) in green. The width of each line represents the magnitude of the coefficients. aDM: anterior default mode network (as opposed to classical (posterior) DM), ATN: attention network, LFP/RFP: left/right frontoparietal network, SUB: subcortical network, MTR: Somatomotor network, AUD: auditory network, CER: cerebellar network, and VIS: visual network.

Additionally, we found the symptom-related DM-FNCs differentiate (FDR < 0.05) between schizophrenia and typical controls in all three AllTPs, EPTs, and EATs models, further supporting their biological relevance. These results suggest that different portions of data, including EATs, may carry unique/additional information about the default mode, and a focus only on EPTs miss this potentially important information.

We also performed additional analysis and evaluated changes within the default mode network. We applied voxel-wise group comparison while including age, gender, site, and mean framewise displacement (mFD) as confound regressions and correcting for multiple comparisons. We found significant differences in the thalamus for the ICA analysis using EATs (Supplementary 10). We did not observe any difference in the EPTs and AllTPs. This may be due to the presence of time-varying spatial connectivity consistent with previous findings (52).

## Discussion and Perspective

The main objective of functional neuroimaging is to relate the measurements of imaging modalities to underlying neural activity and biological variability. Among different imaging modalities, fMRI, which measures the BOLD signal as a proxy for average neural activity, has advantages over others because of its ability to record data from the whole brain, arguably the best trade-off between spatial and temporal resolutions. However, there is still little known about the underlying functional architecture and how neural activity contributes to the spontaneous BOLD signal. To decipher this mystery, the neuroimaging community has dedicated great effort to develop analytical approaches to model spontaneous BOLD signal and interpret the underlying functional patterns.

Building upon the notion of dynamic, ongoing functional interactions across the brain, the dominant category of approaches model intrinsic functional patterns (i.e., functional networks) as continuous entities in time. A major departure from this category seeks to model brain function as a set of discrete (i.e., sparse in time) functional patterns, initially developed based on the hypothesis that spontaneous BOLD signal results from brief and isolated epochs of neural events (8). Because detecting event present time points is the core component and the real differentiator of this category, we used the term “event detection approaches” as the general term to describe this category of approaches. Similar to the main category, event detection studies have presented intriguing findings, suggesting their potential to facilitate understanding of underlying neural activities with great opportunities for both clinical and research settings. This results in a rapidly growing interest in event detection approaches and using only a subset of data (EPTs), instead of all available time points to study brain function.

This study argues that intrinsic networks have an ongoing, continuous presence in BOLD signal, and EPTs do not contain all the information of their corresponding functional patterns. As such, using only EPTs in fMRI studies may result in missing important information, potentially leading to misleading conclusions. The findings of this study support this proposition. Thus, while we emphasize the potential of event detection approaches and advocate for further studies of this category, we call for caution on the assumptions and interpretation of the findings.

We summarize the findings of this study in three main analyses. The first analysis stemmed from the premise that network patterns derived from discrete, temporally sparse events, compared to continuous ongoing interactions between associated regions. We focused on the default mode pattern because it has been repeatedly detected as a dominant global co-activation pattern and arguably most studied in ROI-based event-driven studies. We evaluated the continuous presence of the default mode in spontaneous BOLD signal by investigating if the default mode significantly exists in the time points with the least probability of being the default mode event (EATs), i.e., the time points with amplitudes near to baseline (zero). We tested our proposition using two commonly used default mode nodes in event-based studies, using a node obtained from meta-analysis for the term “default mode” in Neurosynth, and even using the time course of subject-specific default mode obtained using group ICA followed by multi-objective optimization (47). We observed that regardless of choice of node, preprocessing steps (cleaning), analysis (first, second, and higher-order statistics), the default mode patterns strongly present in both activity and connectivity spaces. These findings suggest that the default mode is a continuous phenomenon or at least has a continuous footprint in spontaneous BOLD signal. Therefore, EPTs do not contain the complete information of functional patterns and might be insufficient to model their properties. Considering that the true nature of the underlying neuronal activity is still unknown, focusing only on EPTs may obscure our understanding of brain function. The success of event detection approaches in identifying brain networks and capturing useful information using only EPTs should not be seen as evidence that networks only present or originated from discrete infrequent events (34).

While the main objective of the first analysis was to evaluate the presence of the default mode at EATs of the default mode, the second analysis targeted its EPTs. We were particularly interested in whether spontaneous BOLD signal in these time points is induced by default mode events and merely reflects the default mode or subsists on other functional patterns, obtained from the whole data. ICA analysis shows that large-scale brain networks can be equally obtained from EPTs as well as EATs. The results of these analyses provide evidence that intrinsic networks have an ongoing, continuous presence in the BOLD signal, and they are best modeled as continuous in both space and time, rather than as discrete events. While our supplementary analyses support our findings for the visual and somatomotor network, the findings of this study should be further evaluated for additional networks and spatial scales.

The third analysis was designed to answer whether EPTs can capture a full picture of functional properties or if different portions of time carry unique/complementary information. First, we studied the association between a schizophrenia symptom severity score and the DM-FNC calculated using only EPTs, only EATs, and AllTPs. We observed common patterns such as the contribution of DM-CER connection across all three models; in addition, robust and unique symptom-related links were found for all three scenarios. For instance, unlike the EPT model, the AllTPs model identified the DM-AUD connection to be related to the symptom score suggesting the EPT approach may be missing some information present in the full-time series. Further supporting this, the DM-LFP connection from the EATs shows a robust association with symptoms. At the same time, we advocate for using EPTs to study brain function as they can contain unique information about brain states. We observed that the DM-RFP connection contributes only to the EPT model. These findings are aligned with previous studies showing relationships between symptom severity and functional connectivity of regions from cerebellar, default mode, and left/right frontoparietal networks (53–57). For instance, Brady et al. identified the functional connectivity of the bilateral dorsolateral prefrontal regions (regions in LFP and RFP) with the default mode regions covaried significantly with symptom severity (53). In particular, the functional connectivity of the right dorsolateral prefrontal was associated with negative symptom severity (53). We observed DM-RFP FNC is associated with PANSS total score in the EPTs. The functional connectivity between a default mode node in posterior cingulate cortex and left middle frontal gyrus (regions involved in LFP) and PANSS general symptom is shown to be correlated (55). Similarly, a DM-LFP FNC association with the PANSS total score was found in the EAT analysis. LFP dysfunction has been frequently associated with a range of abnormalities in schizophrenia, correlated with PANSS score, and suggested as a potential endophenotypic marker of schizophrenia (56). The relationship between the PANSS and the default mode has also been reported for within-DM functional connectivity (58). Strikingly for all three models (i.e., AllTPs, EPTs, and EATs), the symptom-related DM-FNCs also differentiated between schizophrenia and typical controls, supporting their potential biological relevance. Building on these findings, we propose different portions of time, to be exact EPTs versus EATs versus AllTPs, carry unique/complementary information, and using EPTs alone may not sufficient to study brain function. Furthermore, because symptom scales like PANSS are the primary clinical tools to assess psychotic behavioral disorders, their functional fingerprints can provide a better understanding of schizophrenia-related brain functional changes. Despite promising findings, the symptom severity scores on the schizophrenia subjects are unfortunately differ between the two samples (PANSS versus BPRS), which we attempt to harmonized them using established a prior conversion algorithm obtained to convert BPRS total scores to PANSS total scores. This, on the other hand, results in very blunt instrument (PANSS total score) compared to the finely tuned tools. We propose that future studies should dedicate more efforts to deciphering behavior-functional imaging relationships and delineating the symptom scale brain functional fingerprint, particularly by leveraging a finer breakdown of severity symptom scores into positive and negative and disorganized symptoms.

In addition to differences in the context of schizophrenia, our analyses illustrated differences in the functional patterns of EPTs and EATs, suggesting they may reflect different states of brain functional architecture. One intriguing finding was observed in the executive control network. The executive control network has consistently been identified as a separate functional network with anticorrelative patterns with the default mode in both time and space. The executive control and default mode were also reported to have opposite responses (activation versus deactivation) during cognitively demanding tasks (59). Similar to previous work, these two appeared as two independent components in our ICA analyses using AllTPs and EPTs. However, for EATs obtained using voxels mainly in the posterior cingulate cortex (i.e., Node_Seed1_, Node_Seed2_, and Node_Meta_), we observed these two networks seem to emerge into one component which contains the same default mode regions as EPTs and AllTPs and many regions of the executive control network (see **Figure 2** and **Figure 3**).

Considering the premise about the central role of the posterior cingulate cortex in coordination between these intrinsic connectivity networks and supporting internally-directed cognition (60), which is important during goal-directed tasks, these moments may reflect information integration between these two networks and modulation of top-down processing (61). This finding is also supported by the recent spatial dynamic observation of dynamic integration and segregation between brain networks (52). Iraji et al. show that intrinsic networks, commonly considered separate entities in previous spatial static studies, transiently merge and separate, reflecting their dynamic segregation and integration (52, 62). Our findings suggest that the default mode and executive networks are fully segregated at EPTs, which is expected as the default mode activity is maximum and main default mode regions are expected to strongly connect to each other compared to their functional connections to other regions. However, in the EATs, the functional connectivity among default mode regions, particularly the posterior cingulate cortex, is lower relative to their connections with other regions and potentially reflects the transfer of information and modulatory interaction between these two momentarily integrated networks. As such, we argue that focusing only on EPTs would be limited to dominant within network dynamics and unveil dynamic segregation patterns, while including EATs would allow us to better capture dynamic integration and spatial fluidity between networks (52).

In addition, existing event-based studies are mainly limited to identifying spatial patterns that resemble large-scale distributed networks obtained in functional connectivity studies. Early event-based studies have used this similarity to support their hypothesis and assert that functional connectivity results from coactivation in EPTs. We argue that while the activation maps of EPTs successfully identify large-scale covarying functional patterns, more spatially local functional connectivity patterns in the data will be missed when using an EPT-type analysis. In other words, the activity patterns cannot explain the functional patterns obtained across multiple spatial scales, such as fine-grained ICNs obtained in higher-order ICA analysis (5, 63). Moreover, studies have shown that functional connectivity can occur in cross-frequency and at different frequencies within a given network (41, 64, 65). As a matter of fact, various types of neural activities with distinct spatial and temporal representation may contribute to the BOLD signal (66, 67).

## Concluding Remarks and Future Considerations

We propose that the BOLD signal can be best modeled as a combination of processes that occur at different spatial and temporal scales. Leveraging different existing models and analytical approaches and developing new ones can help better understand underlying neuronal processes that contribute to the BOLD signal, leading to better insights into brain function and its alterations in various brain conditions. This includes event detection approaches and other single frame-based techniques (use single time points as the elements of analysis). However, one should consider the impact of noise on these approaches considering the signal-to-noise ratio of the BOLD signal.

Furthermore, the key step in event detection approaches is to accurately identifying a functional pattern time series to detect time point. we recommend leveraging data-driven approaches instead of predefined anatomical nodes to more accurately obtain time series and detect EPTs. When a region is a central part of a functional pattern (e.g., the posterior cingulate cortex for the default mode), its high amplitude time points can effectively depict the functional pattern. However, as our results show, the amplitude of a given fixed node does not well represent the activity of the corresponding functional pattern, rather the activity of a given node and its contribution to functional patterns. For instance, while EPTs obtained from a posterior cingulate cortex node extract the default mode, the default mode is also strongly present in its EATs, but the node’s contribution to the default mode is not as strong as the rest of the default mode regions. This, for example, can be seen in the top right panel of **Figure 2**, which shows the default mode network obtained by applying ICA on the EATs of Node_Seed1,_ where the node itself appears as a ‘hole’ and does not have a strong contribution to the default mode. This is expected given the EATs were selected to have minimal relationship to Node_Seed1_, but it is striking that even in this extreme case the rest of the default mode can still be well-estimated.

Related to this, different methods capture different information regarding the association of a region to a functional pattern. Both ASM and FCM (first and second-order statistics) calculate the contributions of regions/voxels to their dominated functional patterns without considering their contribution to other functional patterns. On the other hand, multivariate analyses like ICA calculate the degree of associations while controlling for the effect of other functional patterns in a model. This can result in differences in spatial maps of different functional patterns (e.g., **Figure 2** and **Figure 3**). We suggest that using multivariate data-driven approaches to extract the time course of a functional pattern might better detect its EPTs. This can be done using fully blind approaches like clustering or ICA (4), or hybrid approaches that use spatial constraints to facilitate comparability across analyses while also adapting to the data to ensure functional coherence (68).

Future studies should assess our findings using other brain networks and use other event detection approaches such as using local maxima and minima, deconvolving the hemodynamic response, or using other approaches such as instantaneous functional connectivity (second or higher-order statistics) instead of activity (first-order statistic). Furthermore, future studies should studies how functional connectivity between network being affected by choice of EATs, EPTs, and AllPTs and should develop approaches to assess reproducibility in the context of time-varying changes (69). Finally, future single frame-based studies, including event detection approaches, should also explore the benefits of incorporating spatial dynamics (52, 62, 70).

In sum, our finding suggests that functional networks are best modeled as continuous, evolving temporal patterns with different portions carrying unique/complementary information. This highlights the necessity of utilizing all time points and opens new possibilities to study brain function.

## Acknowledgements

This work was supported by grants from the National Institutes of Health grant numbers 1U24RR021992, 1U24RR025736, R01EB020407, and R01MH118695, and National Science Foundation grant 2112455 to Dr. Vince D. Calhoun and by grants from the VA Merit I01CX000497 program and VA Senior Research Career Award to Dr. Judith M Ford.

